# Evaluating the estimability of within-host population dynamics models

**DOI:** 10.64898/2026.08.21.746183

**Authors:** Madeline Jarvis-Cross, Andrew W. Bateman, Cole B. Brookson, Nicole Mideo, Martin Krkošek

## Abstract

Despite the impacts of within-host disease dynamics on disease outcomes in individual hosts and disease spread among-hosts, generic models of within-host population dynamics have received far less attention than their among-host counterparts. While a number of models have been proposed to explore theoretical eco-evolutionary dynamics, they have yet to be evaluated for estimability, raising questions about their ability to provide reliable inference when confronted with data.

We evaluated the estimability of two generic within-host population dynamics models by assessing: (1) parameter estimation, our ability to recover correct values of model parameters from data, (2) the consequences of mis-assigning the underlying mechanistic model on parameter estimation, and (3) the reproduction of qualitative dynamics, or, our ability to use parameter estimates to reproduce observed dynamical behaviours. In some cases, fitting a mis-matched mechanistic model to time series data produced reasonable parameter estimates that were able to reproduce system dynamics, and that when provided the data-generating model, parameter uncertainty can produce substantial behavioural uncertainty. Our findings highlight the impacts of structural, parametric, and behavioural uncertainty on inference, and demonstrate the value of improving system-specific knowledge to prevent the use of incorrect functional forms and of measuring consequential parameters to improve estimability.

## Introduction

Within-host disease dynamics influence infection intensity, host mortality rates, and among-host transmissibility [1, 2]. Within-host population dynamics models often draw from consumer-resource theory and Lotka-Volterra models [3–7], which have been found to reliably describe the population dynamics of interacting predator and prey species [8], such as lynx-hare [9], herbivore-plant [10, 11], bird-microtine [12], raptor-grouse [13], and wolf-moose systems [14]. Within-host models usually represent a host’s immune system as attacking infecting parasites analogously to how a predator attacks and consumes prey, and include functional responses to describe an immune system’s response to infection intensity [15–19]. For example, as a predator’s consumption of prey items may be mitigated by their attack rate and the time it takes to consume prey items, producing a saturating response to increased prey abundance, observed relationships between the immune system and an invading parasite suggest that similar processes can produce comparable dynamics within hosts [20] (e.g., the activation of the innate immune response and the development and proliferation of the adaptive immune response [19], the time it takes for an immune cell to dispatch a parasite [21, 22]).

This conceptual underpinning has been invoked to build simple models of interactions between known host cells and parasites, and estimate parameter values from model fits (e.g., [23–27]). However, building and testing models of poorly understood within-host disease systems can be particularly difficult due to the fact that it is often unclear which components of the immune system or which immune and infection processes should be modelled or measured. For example, in vertebrates, the innate and adaptive immune systems may interact with infecting parasites differently [26] and the cells that comprise these systems differ in their proliferation and dispatchment of parasites [22]. These uncertainties demonstrate a clear use-case for generic within-host population dynamics models (from which system-specific models may be elaborated). However, despite generic predator–prey models having been fit to data [8], their within-host descendents (e.g., [16, 18, 19, 22]) have yet to be evaluated for estimability, raising questions about their ability to provide reliable inference when confronted with data.

In contrast, at the among-host level, compartmental models broadly describe the movement of a pathogen through a susceptible population [28–30]. SIR (Susceptible → Infected → Removed) models divide the host population among different compartments, and describe their movement from one compartment to another. For example, individuals in the “Susceptible” compartment may come into contact with individuals in the “Infected” compartment, and become infected, moving from the “Susceptible” compartment to the “Infected” compartment. Similarly, individuals in the “Infected” compartment may either recover and gain immunity to future infection or succumb to infection, moving from the “Infected” compartment to the “Removed” compartment. Since its development, the SIR model has been elaborated on to describe various disease systems with more specificity, for example: modelling the spread of SARS-CoV-2 with a Susceptible → Exposed → Infected → Removed framework [31–35]. Importantly, generic and system-specific SIR models have been evaluated as estimable and have been used to estimate key parameters (e.g., transmission rate) and metrics, including the basic reproductive number, *R*_0_, interpreted as the number of secondary infections expected from a single infection within a susceptible population [36] (e.g., [33, 37]).

We evaluated the estimability of two generic within-host population dynamics models to assess the utility of generic within-host population dynamics models in poorly-understood systems. Antia et al. (1994) and Fenton and Perkins (2010) propose within-host population dynamics models within which the interaction between the host’s immune system and an infecting parasite is analogous to the interaction between a predator and its prey [22, 38]. While neither model was originally developed to infer system-specific dynamics, we used these models as starting points to evaluate the utility of generic within-host population dynamics models by assessing: (1) parameter estimation, or, our ability to recover the values of model parameters from data, (2) the consequences of mis-assigning the underlying mechanistic model on parameter estimation, and (3) the faithful reproduction of system dynamics, or, our ability to use parameter estimates to reproduce observed behaviours [39, 40]. We used the two within-host population dynamics models to simulate demographically stochastic infection time series, and backfit the generating models to the time series data. Then, we evaluated the accuracy and precision of parameter estimates, the likelihood of mistaking a mis-assigned underlying model for the data-generating model, and the ability of the fitted model to reproduce the expected qualitative dynamics (e.g., damping oscillations, divergent oscillations, etc.). We identified key parameters whose fixation would improve the estimation of other parameters, and found that uncertainty in parameter estimates made it difficult to reproduce qualitative dynamics, calling into question our ability to predict infection outcomes, and demonstrating the importance of developing system-specific knowledge for specifying functional responses.

## Materials and methods

To assess the estimability of within-host population dynamics models, we used two within-host models to simulate stochastic time series data, and then fitted the generating models to the simulated data in a Bayesian framework. We begin this section by describing Antia et al.’s (1994) and Fenton and Perkins’ (2010) within-host population dynamics models, their parametrisations, and dynamical behaviours. Then, we describe how we generated deterministic and demographically stochastic time series of parasite and immune cell dynamics. Finally, we describe the statistical models we fit to these data in global and autoregressive frameworks, how we assessed model identifiability and goodness of fit, and how we used model outputs to conduct a behavioural uncertainty analysis (Fig. 1).

**Fig 1.**
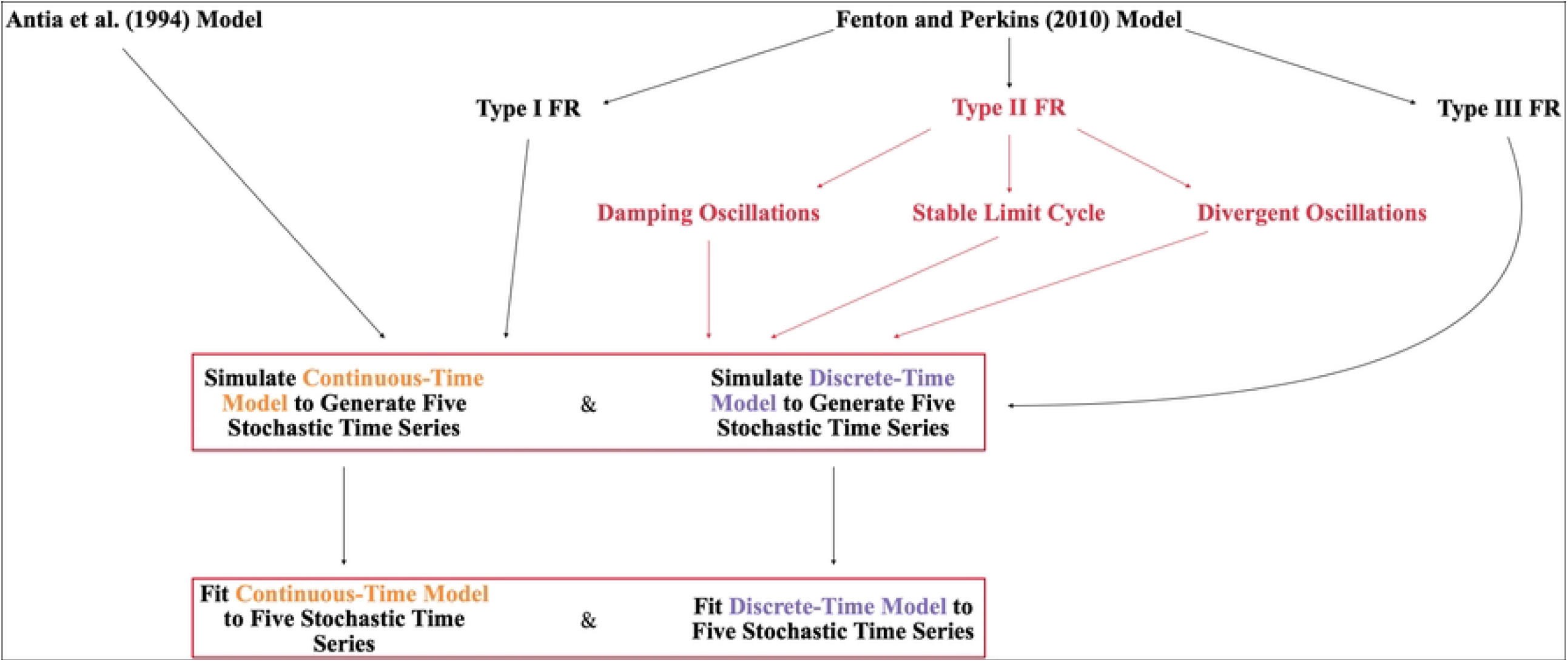
Schematic detailing the simulation experiment. We explore the stability of two models, proposed by Antia et al. (1994) and Fenton and Perkins (2010). We specify three versions of the Fenton and Perkins (2010) model, which are differentiated by the response of the host’s immune system to parasite density. When the host’s immune system has a Type II functional response (FR) to parasite density, different parametrisations of the model produce different dynamical behaviours (highlighted in red). We use continuous-(orange) and discrete-time (purple) implementations of these models to generate stochastic time series data, and fit the generating models to these data.

## Model descriptions

Antia et al.’s (1994) and Fenton and Perkins’ (2010) within-host population dynamics models both describe a microparasite infection, within which the interaction between the host’s immune system and an infecting parasite is analogous to the interaction between a predator and its prey. Both models are biologically and mathematically simple enough as to be broadly applicable to an array of within-host microparasite systems and analytically tractable. Antia et al.’s (1994) model describes the interaction between the host’s immune system and the parasite as short-lived, resulting in either clearance or host death [38]. In contrast, Fenton and Perkins’ (2010) model can be formulated in three different ways to specify the immune system’s response to infection intensity, and describes a sustained infection [22].

### Model (1) from Antia et al. (1994)

Antia et al. (1994)’s model of within-host population dynamics describes a system within which the interaction between an infecting parasite, *P*_*A*_, and the host’s immune system, *I*_*A*_, is analogous to the interaction between a prey and its predator (Eq. (1.1.1) and (1.2.1), Table 1). Per Antia et al. (1994), parasite dynamics are dictated by the parasite replication rate, *r*, and the rate at which parasites are consumed by immune cells, *k*. The model assumes that the replication rate of the host’s immune system is proportional to parasite density at low parasite densities, and saturates at a maximum rate, *ρ*, when parasite densities are high. This formulation allows the host’s immune system to clear or succumb to infection, resulting in recovery or host death, which occurs when parasite density surpasses some threshold, *D* [38]. The dynamics are therefore described by

**Table 1.** Variable and parameter definitions and values for Antia at al. (1994) within-host population dynamics model. *X*^*†*^ denotes an initial condition. Given that Antia et al. (1994) were interested in characterising the behaviour of a generic system rather than representing an empirical system, units are unspecified, but rates are assumed to apply per unit time.

| Symbol | Parameter | Values Stated in Antia et al. (1994) | Values Used to Simulate Continuous- and Discrete-Time Models |
| --- | --- | --- | --- |
| $P_A$ | Parasite abundance | $1^\dagger$ | $1^\dagger$ |
| $I_A$ | Host immune cell abundance | $1^\dagger$ | $1^\dagger$ |
| $r$ | Parasite replication rate | $0.1 - 10$ | $0.2 - 0.25$ |
| $k$ | Rate at which parasites are consumed by immune cells | $10^{-3}$ | $10^{-2}$ |
| $\rho$ | Maximum replication rate of immune cells | $1.0$ | $1.0$ |
| $\phi$ | Parasite density at which replication rate of immune cells is half its maximum | $10^3$ | $10^3$ |
| $D$ | Lethal within-host parasite density | $10^9$ | $10^9$ |

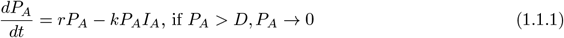

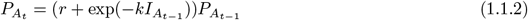

where *ϕ* is the parasite density at which the immune cells replicate at half its maximum. The model can be implemented in discrete-time as follows:

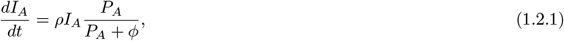

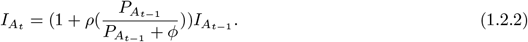

### Model (2) from Fenton and Perkins (2010)

Fenton and Perkins (2010) explore microparasite-host dynamics using three formulations of a Lotka-Volterra model (Eq. (2.1.1) and (2.2.1)). Within each formulation, changes in parasite load,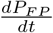, are dictated by the parasite replication rate, *r*, and the rate at which parasites are consumed by immune cells, *I*_*F P*_, which is itself dictated by a functional response (FR), *f* (*P*_*F P*_) [22]. The immune system is stimulated at rate *e* and decays at rate *δ* giving,

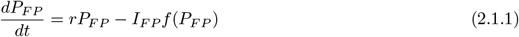

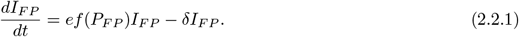

The functional response, *f* (*P*_*F P*_), can be defined as a Type I, Type II, or Type III as follows (Eq. (3.1.1) to (3.3.1), Table 2):

**Table 2.** Variable and parameter definitions and values for Fenton and Perkins (2010) within-host population dynamics model. *X*^*†*^ denotes an initial condition.

| Symbol | Parameter | Values Stated in Fenton and Perkins (2010) | Values Used to Simulate Continuous- and Discrete-Time Models |
| --- | --- | --- | --- |
| $P_{FP}$ | Parasite abundance | $1^\dagger$ | $1 - 80^\dagger$ |
| $I_{FP}$ | Host immune cell abundance | $1^\dagger$ | $1 - 200^\dagger$ |
| $r$ | Parasite replication rate | $0.05 - 0.1$ | $1.5 - 3.0$ |
| $\beta$ | Rate at which parasites are consumed by immune cells | $0.05$ | $0.001 - 0.012$ |
| $h$ | Immune response "handling time" | $0 - 10$ | $0 - 0.2$ |
| $e$ | Immune stimulation rate | $0.1$ | $0.2 - 0.9$ |
| $\delta$ | Decay rate of immune system | $0.01$ | $0.2 - 0.5$ |
| $b$ | Innate immune system | $-$ | $35 - 200$ |

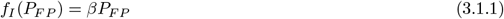

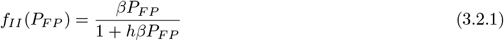

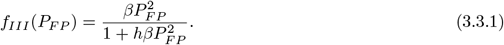

A Type I response describes the relationship between parasite density and the elimination of parasites by the immune system, *β*, as linear, while a Type II response as saturating, and a Type III response as sigmoidal to represent a lag in the immune response a low parasite densities and saturation at high parasite densities. It has been posited that a Type I response may describe a system within which immune memory promotes a rapid immune response to a familiar parasite [41], while a Type II response may be representative of systems within which the handling time, *h*, required for a macrophage or phagocytic cell to dispense with an infected cell or extracellular parasite is considerable [19, 20]. Type III responses may be representative of the immune system’s response to a novel parasite of which it has no memory [19, 21, 22, 42, 43]. Both Type II and III functional responses cause the immune response to saturate at high parasite densities, suggesting that the immune response is limited by the host’s resources, and cannot grow indefinitely.

In contrast to Antia et al.’s (1994) model, which captures an acute infection, all three formulations of Fenton and Perkins’ (2010) model describe a sustained infection. Additionally, different formulations and parametrisations of the model produce different dynamical behaviours (Fig. 1, Table 3; see Fenton and Perkins, 2010 Supplementary Material for stability analyses). Per Fenton and Perkins (2010), when the immune system exhibits a Type I response to infection, the model produces a stable limit cycle. When the immune system exhibits a Type II response to infection, different parametrisations of the model produce a stable limit cycle or divergent oscillations [22]. When the immune system exhibits a Type III response to infection, different parametrisations of the model produce damping oscillations or a stable limit cycle [22].

**Table 3.** Focal models, formulations, and the dynamical behaviours they produce. The downward arrow indicates that the formulation of the Antia et al. (1994) model dictates an immune response that shares features with Type I and Type II functional responses.

|  | Antia et al. (1994) | Fenton and Perkins (2010) | Modified Fenton and Perkins (2010) |
| --- | --- | --- | --- |
| Type I | Exponential growth to extirpation | Stable limit cycle | Damping oscillations |
| Type II | ↓ | Stable limit cycle<br>Divergent oscillations | Damping oscillations<br>Stable limit cycle<br>Divergent oscillations |
| Type III | — | Damping oscillations<br>Stable limit cycle | Damping oscillations |

While this model accounts for the immune system’s response to infection, we adapted the model to include constitutive production of immune cells [19, 44], denoted by *b* (Eq. (4.1.1) and (4.2.1), Table 2),

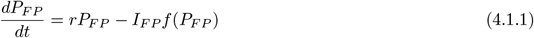

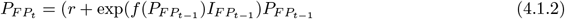

The functional response, *f* (*P*_*F P*_), can again be defined as a Type I, Type II, or Type III (Eq. (3.1.1) to (3.3.1)). As above, different formulations and parametrisations of the model produce different dynamical behaviours (Table 3). Via numerical simulation, we determined that when the immune system exhibits a Type I response to infection, the model produces damping oscillations. When the immune system exhibits a Type II response to infection, different parametrisations of the model produce damping oscillations, a stable limit cycle, or divergent oscillations. When the immune system exhibits a Type III response to infection, the model produces damping oscillations.

The model can be implemented in discrete-time as follows:

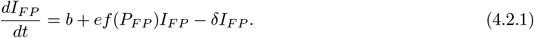

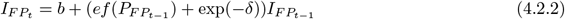

To restrict the scope of the work, maintain a range of dynamical behaviours, and enable comparison between three behaviours produced by the Type II model, we will use the Antia et al. (1994) and Modified Fenton and Perkins (2010) models in all analyses moving forward.

### Model simulations

To generate time series data, we simulated each model forward in continuous and discrete time. In both cases, we simulated deterministic and demographically stochastic implementations of the models, and simulated the models for long enough to observe the extirpation of the parasite (Model (1)) or characterise dynamical behaviours (Model (2)). Demographic stochasticity was introduced via the Gillespie algorithm in continuous time [45] and by writing replication terms as Poisson processes and mortality terms as binomial processes in discrete time. We simulated five time series in continuous time (Fig. 2), and five time series in discrete time (Fig. 3) to account for the impacts of stochastic variation on our findings. Working with five time series enabled us to characterise a range of outcomes within a manageable amount of computational time.

**Fig 2.**
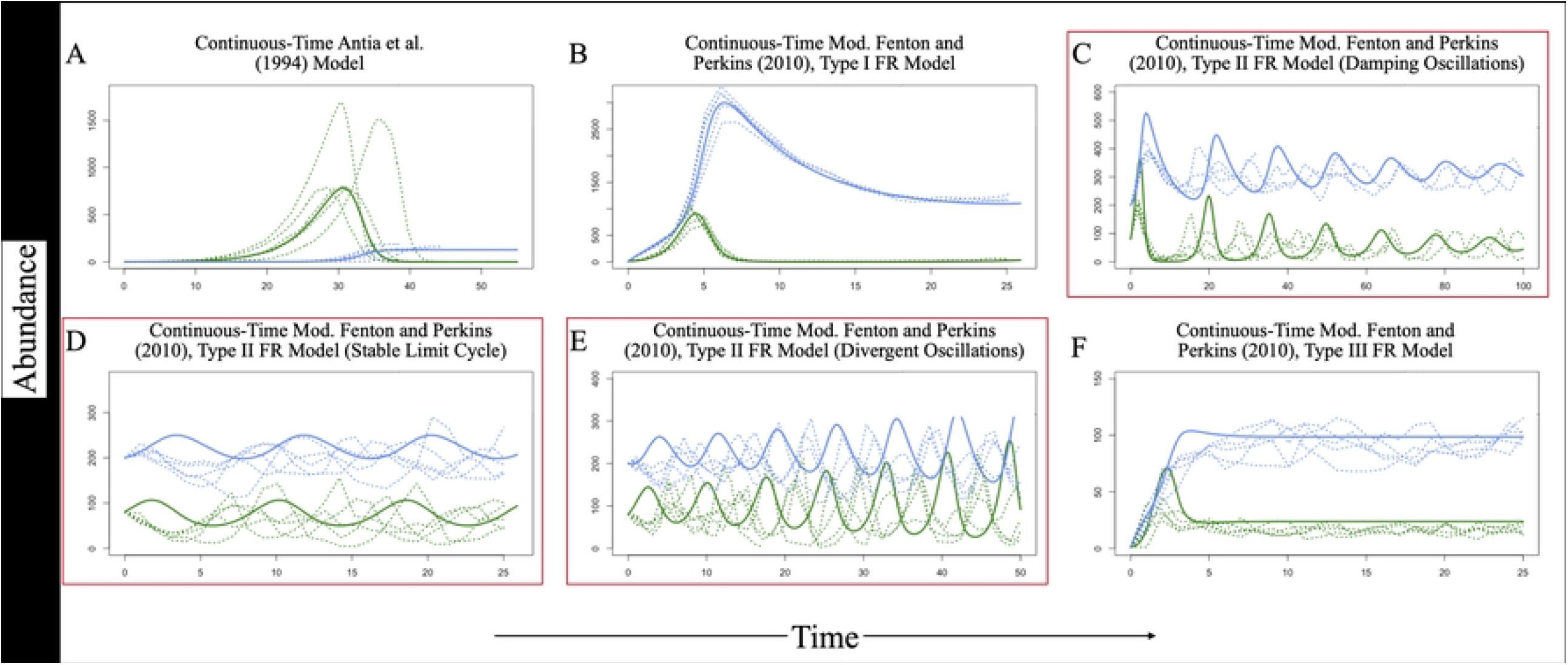
Deterministic and stochastic time series of host immune cell abundances and parasite loads, per each continuous-time model and parametrisation. In each plot, host immune cell abundances are plotted in blue, while parasite loads are plotted in green. Deterministic time series are denoted by solid lines, while stochastic time series are denoted by dotted lines. Red text denotes three parametrisations of the Modified Fenton and Perkins (2010) model with a Type II functional response.

**Fig 3.**
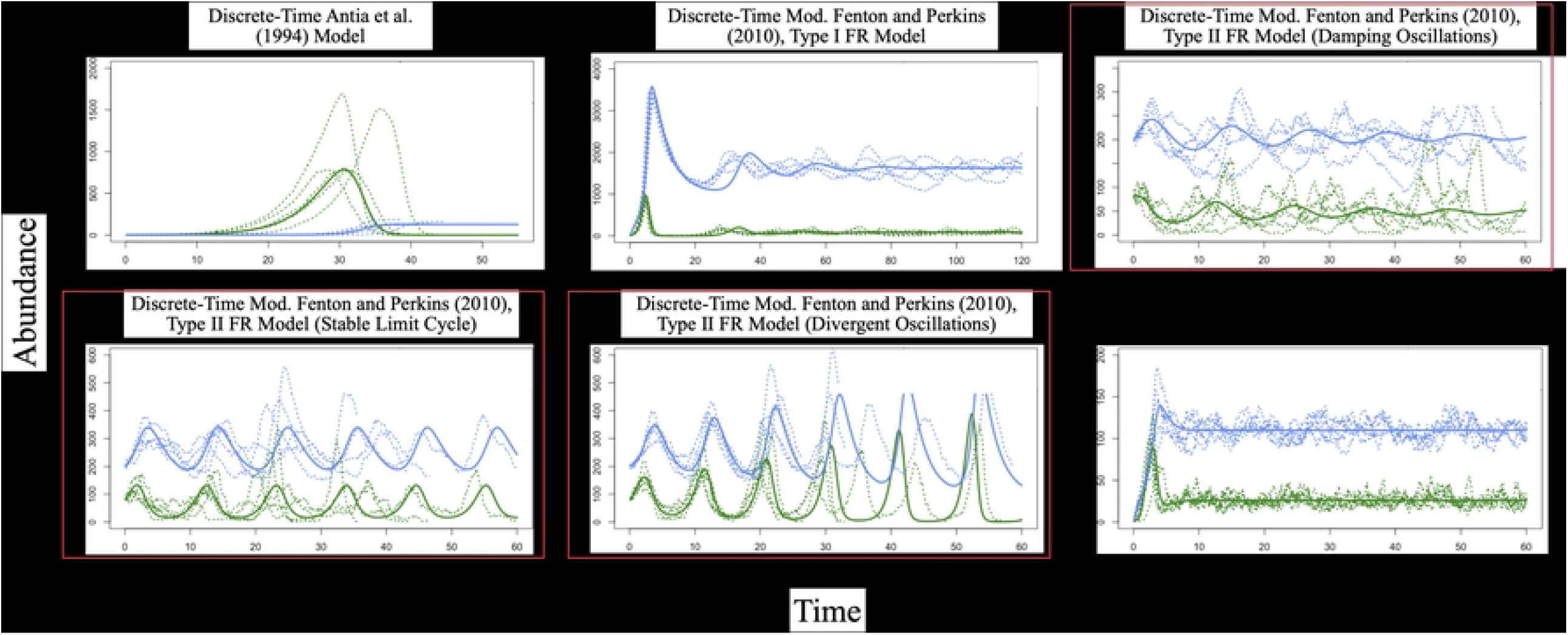
Deterministic and stochastic time series of host immune cell abundances and parasite loads, per each discrete-time model and parametrisation. In each plot, host immune cell abundances are plotted in blue, while parasite loads are plotted in green. Deterministic time series are denoted by solid lines, while stochastic time series are denoted by dotted lines. Red text denotes three parametrisations of the Modified Fenton and Perkins (2010) model with a Type II functional response.

### Model fitting

We fit the models to the simulated data using global and autoregressive (AR-1) frameworks. Within a global framework, we use the model to predict the full time series. Within an autoregressive (AR-1) framework, we use the model to predict the next data point, conditional on the current datapoint, sequentially through the time series. We fit each model in global and autoregressive frameworks to compare each statistical model’s ability to fit generating models to oscillatory data.

To assess the estimability of each generating model, we fit each generating model to its corresponding time series. As an example, we fit the Modified Fenton and Perkins (MFP) (2010), Type II model to five time series generated by each of three different parametrisations of the MFP, Type II model. When fitting the MFP models to time series, we began by estimating all five to six parameters. After encountering difficulty, we provided the fitting model with a single parameter value, and then, with combinations of two parameter values (10 to 15 unique combinations) to identify key parameters whose fixation improved the estimation of other parameters, and the reproduction of system dynamics.

To assess the consequences of mis-assigning the underlying mechanistic model on parameter estimation, we fit each mechanistic model to “mismatched” time series data. As an example, we fit the MFP, Type I model to stochastic time series generated by the Antia et al. model, the MFP, Type II model, and the MFP, Type III model.

We assessed model convergence using two metrics: Effective Sample Size (ESS) and 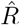 [46]. ESS is a measure of the degree to which within-chain autocorrelation decreases confidence in model estimates [47], and 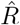 is a metric that compares between- and within-chain variance. When Markov chains mix well and sample from similar distributions, 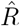 will approach one [46]. While model convergence indicates that the sampler is drawing from a stationary target distribution [48], convergence itself does not guarantee that a target distribution is a reliable reflection of the “true” distribution of the data.

### Global model-fitting framework

We fit each continuous-time generating model (Antia et al.; MFP, Type I; MFP, Type II; MFP, Type III) to stochastic time series using ‘rstan’ [49]. We defined all parameters as real and non-negative, and specified the use of a stiff ordinary differential equation (ODE) solver and the backward differentiation formula (BDF) method to accommodate difficult numerical solutions [50, 51]. We then sampled from the posterior using the No-U-Turn Sampler (NUTS; [52]). We employed weakly informative priors and defined population abundances as lognormally distributed. (To read the models in their entirety, please refer to the [online repository](https://github.com/MadelineJC/Estimability_Repo).) We sampled across four chains for two thousand iterations.

### Autoregressive model-fitting framework

We fit each discretised generating model (Antia et al.; MFP, Type I; MFP, Type II; MFP, Type III) to stochastic time series using ‘rstan’ [49]. We defined all parameters as real and non-negative and added an additional upper-bound of one to proportional parameters. Within the likelihood function, we generated estimates as conditional on the previous time step. As above, we employed weakly informative priors and defined estimated population abundances as normally distributed. (To read the models in their entirety, please refer to the [online repository](https://github.com/MadelineJC/Estimability_Repo).) We sampled across four chains for ten thousand iterations.

## Behavioural uncertainty analysis

As stated above, different parametrisations of the MFP, Type II model give rise to different dynamical behaviours (damping oscillations, stable limit cycle, or divergent oscillations). We conducted a behavioural uncertainty analysis to assess the effects of parameter uncertainty on behavioural uncertainty, or, our ability to correctly characterise the dynamical behaviour of the generating model. Per parametrisation, we drew randomly sampled parameter combinations from the posterior distribution, and simulated the deterministic implementation of the MFP, Type II model forward in time. We then quantified the proportion of simulated time series that exhibited the expected dynamical behaviour by comparing the amplitudes of oscillations at the beginning of a time series to those at the end. We interpreted decreasing amplitudes as damping oscillations, consistent amplitudes as stable limit cycles, and increasing amplitudes as divergent oscillations.

## Results

### Model fitting

#### Global model-fitting framework

Fitting the Antia et al. model to matching time series resulted in convergence and produced reasonable parameter estimates that reproduced system dynamics in five of five cases (S1 Fig, S2 Fig, S3 Table). While fitting the Antia et al. model to mis-matched time series sometimes resulted in convergence, estimated abundances failed to reproduce system dynamics (Table 4).

**Table 4.** Summary of global model fits. Each row indicates a model being fit to data, while each column indicates the model used to generate a set of five stochastic time series (“DampO” → “Damping Oscillations”, “SLC” → “Stable Limit Cycle”, “DivO” → “Divergent Oscillations”). As such, each cell summarises the outcome of fitting a given model to a given set of time series. The number indicates how often the model converged (e.g., “4” indicates that the model converged when fit to four of five time series). Bolded numbers indicate that the generating model and time series “match”. Green cells indicate that estimated abundances reproduced system dynamics, while red cells indicate that estimated abundances failed to reproduce system dynamics. Yellow cells indicate that estimated abundances reproduced system dynamics in some cases.

|  |  | Data |  |  |  |  |  |
| --- | --- | --- | --- | --- | --- | --- | --- |
|  |  | Antia et al. | MFP, Type I | MFP, Type II (DampO) | MFP, Type II (SLC) | MFP, Type II (DivO) | MFP, Type III |
| Model | Antia et al. | **5** | 2 | 1 | 4 | 3 | 1 |
|  | MFP, Type I | 5 | **4** | 5 | 4 | 4 | 5 |
|  | MFP, Type II | 4 | 3 | **2** | **3** | **2** | 2 |
|  | MFP, Type III | 3 | 0 | 0 | 0 | 0 | **4** |

Fitting the MFP, Type I model to five matching time series resulted in convergence and produced reasonable parameter estimates that reproduced system dynamics in four of five cases (S1 Fig, S2 Fig, S4 Table). In general, fitting the MFP, Type II model to matching time series proved difficult. Fitting the MFP, Type II model to time series generated by the damping oscillations, stable limit cycle, and divergent oscillations parametrisations of the model resulted in convergence and produced reasonable parameter estimates in two, three, and two of five cases, respectively (S1 Fig, S5 Table, S6 Table, S7 Table). Notably, fitting the model to data generated by the stable limit cycle parametrisation of the model produced divergent oscillations, rather than a stable limit cycle, in two of three cases (S2 Fig). In contrast, fitting the MFP, Type III model to five matching time series resulted in convergence and produced reasonable parameter estimates that reproduced system dynamics in four of five cases (S1 Fig, S2 Fig, S8 Table).

Surprisingly, in most cases, fitting the MFP, Type I model to mis-matched time series resulted in convergence and produced estimates that reproduced system dynamics (Table 4). Fitting the MFP, Type II model to data generated by the Antia et al. and MFP, Type III models resulted in convergence in four and two of five cases, respectively (Table 4). In these cases, estimated abundances reproduced system dynamics (Table 4). Fitting the MFP, Type II model to data generated by the MFP, Type I model resulted in convergence in three of five cases (Table 4). In these cases, estimated abundances did not reproduce system dynamics (Table 4). While fitting the MFP, Type III model to data generated by the Antia et al. model resulted in convergence in three of five cases (Table 4), estimated abundances only reproduced system dynamics in two of these cases (Table 4).

After providing the true value of one parameter, the rate at which parasites are consumed by immune cells (*β*), fitting the MFP, Type II and Type III models to matching time series data resulted in convergence and produced reasonable parameter estimates in five, four, three, and five of five cases (S3 Fig, S9 Table, S10 Table, S11 Table, S12 Table). In these cases, model estimates reproduced system dynamics (S4 Fig).

#### Autoregressive model-fitting framework

Fitting the Antia et al. model to matching time series resulted in convergence and produced reasonable parameter estimates that reproduced system dynamics in five of five cases (S1 Fig, S5 Fig, S13 Table). As above, while fitting the Antia et al. model to mis-matched time series sometimes resulted in model convergence, estimated abundances failed to reproduce system dynamics (Table 5).

**Table 5.** Summary of autoregressive model fits. Each row indicates a model being fit to data, while each column indicates the model used to generate a set of five stochastic time series (“DampO” → “Damping Oscillations”, “SLC” → “Stable Limit Cycle”, “DivO” → “Divergent Oscillations”). As such, each cell summarises the outcome of fitting a given model to a given set of time series. The number indicates how often the model converged (e.g., “4” indicates that the model converged when fit to four of five time series). Bolded numbers indicate that the generating model and time series “match”. Green cells indicate that parameter estimates reproduced system dynamics, while red cells indicate that parameter estimates failed to reproduce system dynamics. Yellow cells indicate that parameter estimates reproduced system dynamics in some cases.

|  |  | Data |  |  |  |  |  |
| --- | --- | --- | --- | --- | --- | --- | --- |
|  |  | Antia et al. | MFP, Type I | MFP, Type II (DampO) | MFP, Type II (SLC) | MFP, Type II (DivO) | MFP, Type III |
| Model | Antia et al. | 5 | 5 | 1 | 1 | 0 | 3 |
|  | MFP, Type I | 3 | 5 | 0 | 0 | 0 | 0 |
|  | MFP, Type II | 5 | 0 | 5 | 5 | 5 | 5 |
|  | MFP, Type III | 5 | 0 | 5 | 5 | 5 | 5 |

Fitting the MFP, Type I model to matching time series resulted in convergence and produced reasonable parameter estimates that reproduced system dynamics in five of five cases (S1 Fig, S5 Fig, S14 Table). In general, fitting the MFP, Type II model to matching time series in an autoregressive framework proved straightforward. Fitting the MFP, Type II model to time series generated by the damping oscillations, stable limit cycle, and divergent oscillations parametrisations of the model resulted in convergence and produced reasonable parameter estimates in all cases (S1 Fig, S15 Table, S16 Table, S17 Table). Notably, resulting parameter estimates did not always reproduce expected dynamics (S5 Fig). Fitting the model to data generated by the stable limit cycle parametrisation of the model produced damping oscillations in two cases, and fitting the model to data generated by the divergent oscillations parametrisation of the model produced a stable limit cycle in one case (S5 Fig). As above, fitting the MFP, Type III model to matching time series resulted in convergence and produced reasonable parameter estimates that reproduced system dynamics in five of five cases (S1 Fig, S5 Fig, S18 Table).

In general, while fitting to mis-matched time series often resulted in convergence, parameter estimates were unable to reproduce system dynamics (Table 5).

### Behavioural uncertainty analysis

In general, our analysis revealed that uncertainty around estimated parameters could lead one to draw incorrect conclusions regarding the dynamical behaviour of a system. We found that this risk is more pronounced when fitting in an autoregressive framework, but that risk is highest when we expect to observe a stable limit cycle.

#### Global model-fitting framework

When fitting the MFP, Type II model to stochastic time series generated by all three parametrisations of the model in a global framework, we observed overlaps between the posterior distributions of *β, h, e*, and *δ* (S6 Fig). Small differences in the values of these parameters can change the dynamical behaviour of the system.

After using estimated parameters to simulate the deterministic implementation of the MFP, Type II model forward in time, we found that the resulting time series exhibited damping oscillations, stable limit cycles, and divergent oscillations in 99%, 40%, and 90% of expected cases, respectively (Fig. 4).

**Fig 4.**
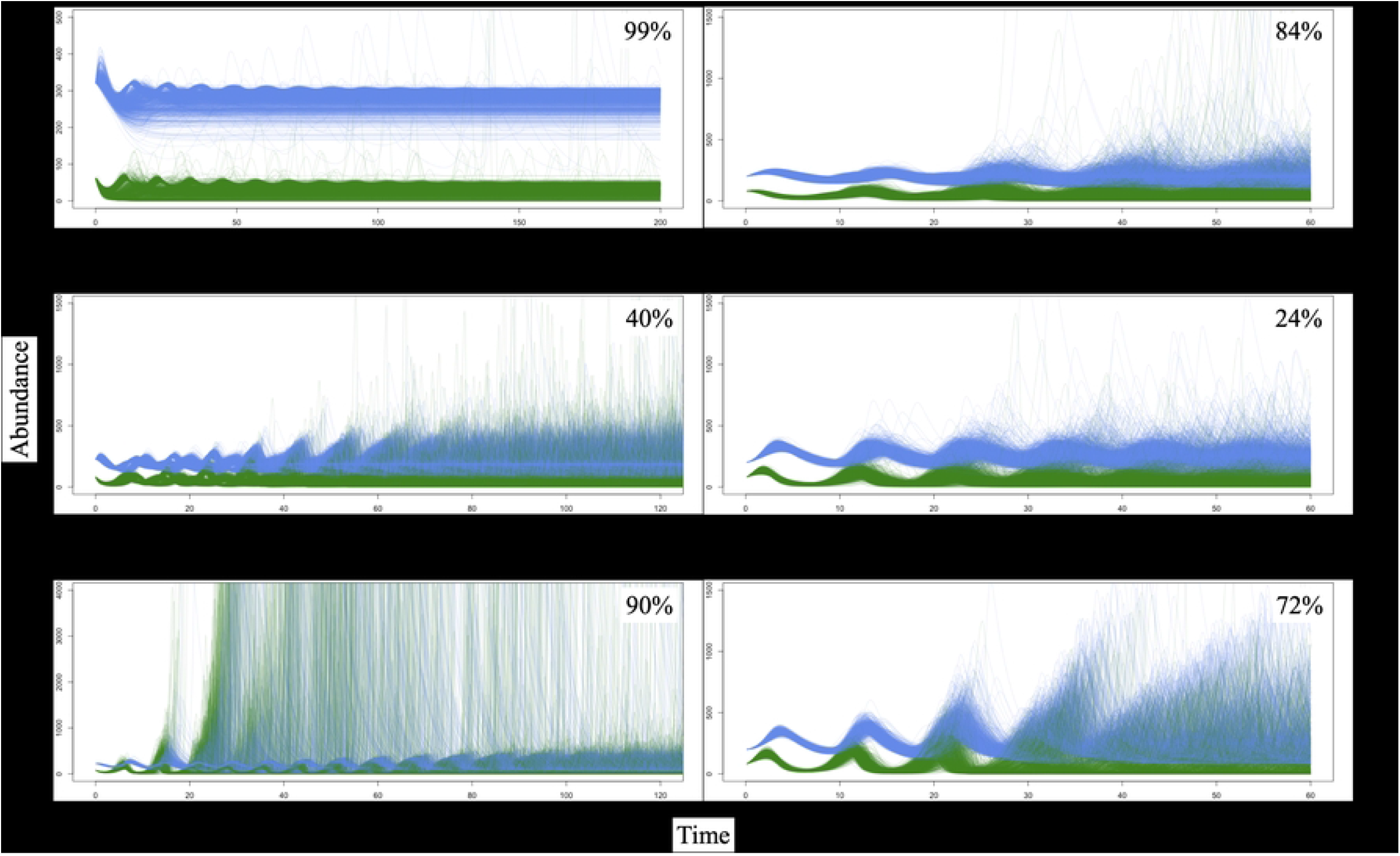
Results of behavioural uncertainty analysis. The MFP, Type II model can produce damping oscillations, a stable limit cycle, or divergent oscillations. After fitting the MFP, Type II model FR model to stochastic time series generated by all three parametrisations of the model, we randomly sampled parameter combinations from the resulting posterior distributions, and simulated the deterministic implementation of the model forward in time. Each plot shows a subset of the resulting time series. Parasite population abundances are shown in green, while immune cell abundances are shown in blue. The value in the top right denotes the percentage of time series that exhibited the expected dynamical behaviour.

#### Autoregressive model-fitting framework

When fitting the MFP, Type II model to stochastic time series generated by all three parametrisations of the model in an autoregressive framework, we observed overlaps between the posterior distributions of *r* and *δ* (S7 Fig). Again, small differences in the values of these parameters can change the dynamical behaviour of the system.

After using estimated parameters to simulate the deterministic implementation of the MFP, Type II model forward in time, we found that the resulting time series exhibited damping oscillations, stable limit cycles, and divergent oscillations in 84%, 24%, and 72% of expected cases, respectively (Fig. 4).

## Discussion

We found that two generic models of within-host pathogen dynamics [22, 38] exhibit a number of estimability issues. Often, fitting a generating model to “matching” time series data produced parameter estimates that were misaligned with data-generating values. However, we identified parameters that, when measured and fixed, improved estimation. In some cases, fitting mis-matched models to time series data produced reasonable parameter estimates that were able to reproduce system dynamics, suggesting that when a system is poorly understood, the risk of mistaking a mis-assigned model for the data-generating model and drawing ill-conceived conclusions about the expected behaviour of a system and an infection’s trajectory might be considerable. Relatedly, we found that uncertainty in parameter estimation made it difficult to reproduce system dynamics, limiting our ability to describe infection dynamics, calling into question our ability to predict infection outcomes from fitted models, and demonstrating the value of measuring consequential parameters to improve estimability.

When fitting mechanistic models to matching stochastic time series data, we found that fitting in a global framework resulted in convergence and in the reproduction of system dynamics less often (4), when compared to fitting in an autoregressive framework (5). This difference was most pronounced when fitting the MFP model to matching data, suggesting that it might be more difficult to infer underlying parameters and biological processes from models that produce an array of dynamical behaviours. Notably, we found that fitting the MFP models to mismatching data in a global framework resulted in convergence and in the reproduction of system dynamics just as often as fitting to matching data did (4). Fitting the MFP, Type I and Type II models to matching stochastic time series in a global framework and providing the true value of one parameter, the rate at which parasites are consumed by immune cells (*β*), more reliably resulted in convergence and in the reproduction of system dynamics. These findings suggest that measuring consequential parameters, like the rate at which parasites are consumed by immune cells or the immune response “handling time” [24, 53, 54], may constrain and improve the estimation of generic within-host population dynamics models [19, 55].

We found that fitting mis-matched mechanistic models to stochastic time series data in a global framework more often resulted in convergence and in the reproduction of system dynamics (4), when compared to fitting in an autoregressive framework (5). Again, this difference was most pronounced when fitting the MFP model to mismatched data, suggesting that more complex models that produce an array of dynamical behaviours may be easier to mistake for the data-generating model. We found that fitting mechanistic models that produced damping oscillations (MFP, Type I and Type II) to mismatched data that exhibited damping oscillations (as generated by MFP, Type I, Type II, and Type III) did not result in convergence more often than fitting mechanistic models to data exhibiting stable limit cycles or divergent oscillations did (4). While deterministic implementations of the MFP models produce damping oscillations, stable limit cycles, and divergent oscillations, stochastic implementations closely resemble one another, suggesting that it may be particularly difficult to differentiate between candidate models when time series data are oscillatory. This problem may be magnified when population dynamics are strongly influenced by stochasticity, or when observations are limited and fail to capture distinguishing behavioural features [56, 57]. Mistaking a mis-assigned model for the data-generating model might lead one to draw ill-conceived conclusions about the expected behaviour of a system and an infection’s trajectory that shape monitoring and treatment regimes and affect infection outcomes. Together, these findings suggest that assessing candidate models of within-host population dynamics by fitting them to data in a global framework and evaluating their ability to reproduce system dynamics may not be sufficient to identify the data-generating model. As such, instead of comparing a wide array of candidate models, we may be better served by improving our understanding of within-host systems to guide the construction of detailed and system-specific candidate models [58].

Similarly, in consumer-resource theory, empirical studies suggest that we may be able to use life history traits to characterise a consumer’s response to prey density. For example, it has been posited that herbivores [10] and specialist predators may be more likely to exhibit a Type II response to prey density, while generalist predators may be more likely to exhibit a Type III response to prey density [59–61]. While it has been suggested that immune memory and handling time may inform our understanding of the immune system’s response to parasite density [19, 22, 41], such relationships have yet to be empirically characterized. As such, determining which life history traits or types of interactions characterise an immune system’s response to parasite density may also help guide the construction of detailed and system-specific candidate models.

In contrast, the behavioural uncertainty analysis revealed that fitting the MFP, Type II model to stochastic time series data in a global framework resulted in less behavioural uncertainty, when compared to fitting in an autoregressive framework. When fitting the MFP, Type II model to stochastic time series data in global and autoregressive framework, we were able to recover damping oscillations, a stable limit cycle, and divergent oscillations in 84% – 99%, 24% – 40%, and 72% – 90% of posterior draws, respectively (Fig. 4). These findings suggest that the analysis of highly sensitive nonequilibrium within-host disease systems would benefit from the measurement of key parameters to minimise uncertainty in the posterior and behavioural uncertainty. The ways in which one approaches parameter or behavioural uncertainty should also be guided by study objectives. For example, limited parameter uncertainty may not be a major concern if one is focused on exploring or describing observed patterns, or inference [58]. Parameter, and thus, behavioural uncertainty may become major concerns if one is focused on prediction [58, 62]. As our simulated time series and findings illustrate, stochastic processes can obscure differences between dynamical systems, and parameter uncertainty can lead to behavioural uncertainty, which can decrease predictive accuracy.

With reference to fitting mechanistic models to time series in global and autoregressive frameworks, fitting the MFP models to stochastic time series data in a global framework resulted in more model uncertainty, when compared to fitting in an autoregressive framework (4, 5). We suspect that this finding is a product of the autoregressive model being able to “follow” the oscillatory behaviour of the system from one time step to the next, minimizing the impacts of model expectations being out of phase with stochastic time series data. As such, our findings suggest that when candidate models or time series data can produce or exhibit complex dynamics, fitting within an autoregressive framework may yield more reliable insights.

Alternatively, one may address these issues by considering an alternative approach. As an example, Wood (2010) proposed reducing time series to phase-insensitive summary statistics that describe local dynamical behaviours, and using these statistics to assess model fit [63]. Separately, fitting the MFP, Type II model to stochastic time series data in a global framework resulted in less behavioural uncertainty, when compared to fitting in an autoregressive framework (Fig. 4). Our findings highlight the benefits of a Bayesian approach, as using posterior probability distributions to quantify parameter uncertainty allowed us to explore the effects of uncertainty on inference and on reproducing system dynamics, and identify circumstances under which model validation would be difficult [62, 64].

Here, we used simulated data to assess the estimability of two within-host population dynamics models with respect to parameter uncertainty, model mis-specification and behavioural uncertainty. Using simulated data allowed us to guarantee the identity of the data-generating model, compare parameter estimates to true values, and assess reproduced time series against the deterministic dynamics of each system. However, to validate within-host population dynamics models (i.e., assess if a model is acceptable for some intended use (here, inferring parameter values and system dynamics); [65]), we must confront these models with empirical data. Notably, given the completeness of simulated time series and the absence of observational and experimental error, our findings likely constitute a best-case-scenario. As such, we expect that introducing empirical data will magnify the complicating impacts of model and behavioural uncertainty on the validation of non-specific within-host population dynamics models.

Our findings demonstrate the ease with which one might mistake a mis-assigned model for the data-generating model, and as such, raise questions about our ability to identify a suitable model via model selection. While nearly ubiquitous in the literature, information criteria have been criticised for using within-sample data to approximate out-of-sample predictive accuracy [66] and implicitly assuming that a system is linear and stationary. Simulation experiments demonstrate that conventional model selection can be unreliable, especially when systems exhibit non-linear and non-stationary (e.g., time-varying, transient) dynamics [67–70]. The estimability issues we have identified would not be well-addressed by conventional model selection approaches, and thus highlight the value of considering biological mechanisms *a priori*.

Despite evidence that within-host dynamics influence disease outcomes in individual hosts and among-host dynamics [1, 2], non-specific within-host population dynamics models have received far less attention than their among-host counterparts. While a number of generic within-host population dynamics models have been developed to explore theoretical interactions between host immune systems and parasites (e.g., [18, 19, 22, 38]), such models have yet to be evaluated for estimability, raising questions about their ability to provide reliable inference when confronted with data. We used two of these models as a starting point to explore the utility of generic within-host population dynamics models, and found that mis-assigned models were easily mistaken for the data-generating model, and that parameter uncertainty begot considerable behavioural uncertainty, raising doubts about the estimability of generic within-host population dynamics models. Our findings underscore the value of developing system-specific knowledge (e.g., measuring consequential parameters) to prevent the mis-assignment of functional forms and to improve parameter estimation and decrease behavioural uncertainty.

## Supporting information

**S1 Table. Parameter values used to generate data to fit continuous- and discrete-time Antia et al. models**.

**S2 Table. Parameter values used to generate data to fit continuous- and discrete-time Modified Fenton and Perkins (MFP) models**.

**S1 Fig. Posterior probability density plots**. After fitting generating models to “matching” stochastic time series, we compared parameter estimates between fits to different time series. Different columns denote different fitting model fitting frameworks, while labels on the left denote the generating model and time series. Each plot shows four to six subplots (one for each free parameter). Each subplot contains five posterior distributions: one for each stochastic time series the generating model was fit to. Vertical lines denote true values.

**S3 Table. Mean parameter estimates and credible intervals resulting from fitting the Antia et al. model to five stochastic time series generated by the Antia et al. model in a global framework**.

**S4 Table. Mean parameter estimates and credible intervals resulting from fitting the MFP, Type I model to five stochastic time series generated by the MFP, Type I model in a global framework**. Grey highlights denote cases in which the model did not converge. Grey colouring denotes cases of non-convergence.

**S5 Table. Mean parameter estimates and credible intervals resulting from fitting the MFP, Type II model to five stochastic time series generated by the damping oscillations parametrisation MFP, Type II model in a global framework**. Grey colouring denotes cases of non-convergence.

**S6 Table. Mean parameter estimates and credible intervals resulting from fitting the MFP, Type II model to five stochastic time series generated by the stable limit cycle parametrisation MFP, Type II model in a global framework**. Grey colouring denotes cases of non-convergence.

**S7 Table. Mean parameter estimates and credible intervals resulting from fitting the MFP, Type II model to five stochastic time series generated by the divergent oscillations parametrisation MFP, Type II model in a global framework**. Grey colouring denotes cases of non-convergence.

**S8 Table. Mean parameter estimates and credible intervals resulting from fitting the MFP, Type II model to five stochastic time series generated by the divergent oscillations parametrisation MFP, Type II model in a global framework**. Grey colouring denotes cases of non-convergence.

**S2 Fig. Stochastic time series and estimated population abundances over time**. After fitting generating models to “matching” stochastic time series in a global framework, we compared the stochastic time series to estimated population abundances. Each plot shows stochastic time series in solid lines, and estimated population abundances in dashed lines. Parasite population abundances are shown in green, while immune cell abundances are shown in blue. Labels on the left denote the generating model and time series. Cases in which the statistical model failed to converge are shown in black and white. Cases in which the behaviour of the stochastic time series appears to be inconsistent with the estimated population abundances are highlighted by red boxes.

**S3 Fig. Posterior probability density plots**. After fitting the MFP, Type II model to “matching” stochastic time series in a global framework and providing the true value of *β*, we compared parameter estimates between fits to different time series. Each plot shows five subplots (one for each free parameter). Each subplot contains five posterior distributions: one for each stochastic time series the generating model was fit to. Vertical lines denote true values.

**S9 Table. Mean parameter estimates and credible intervals resulting from fitting the MFP, Type II model to five stochastic time series generated by the damping oscillations parametrisation MFP, Type II model in a global framework, and providing the true value of** *β*.

**S10 Table. Mean parameter estimates and credible intervals resulting from fitting the MFP, Type II model to five stochastic time series generated by the stable limit cycle parametrisation MFP, Type II model in a global framework, and providing the true value of** *β*.

**S11 Table. Mean parameter estimates and credible intervals resulting from fitting the MFP, Type II model to five stochastic time series generated by the divergent oscillations parametrisation MFP, Type II model in a global framework, and providing the true value of** *β*. Grey colouring denotes cases of non-convergence.

**S12 Table. Mean parameter estimates and credible intervals resulting from fitting the MFP, Type III model to five stochastic time series generated by the MFP, Type III model in a global framework, and providing the true value of** *β*.

**S4 Fig. Stochastic time series and estimated population abundances over time**. After fitting generating models to “matching” stochastic time series in a global framework and providing the true value of, we compared the stochastic time series to estimated population abundances. Each plot shows stochastic time series in solid lines, and estimated population abundances in dashed lines. Parasite population abundances are shown in green, while immune cell abundances are shown in blue. Labels on the left denote the generating model and time series. Cases in which the statistical model failed to converge are shown in black and white.

**S13 Table. Mean parameter estimates and credible intervals resulting from fitting the Antia et al. model to five stochastic time series generated by the Antia et al. model in an autoregressive framework**.

**S14 Table. Mean parameter estimates and credible intervals resulting from fitting the MFP, Type I model to five stochastic time series generated by the MFP, Type I model in an autoregressive framework**.

**S15 Table. Mean parameter estimates and credible intervals resulting from fitting the MFP, Type II model to five stochastic time series generated by the damping oscillations parametrisation MFP, Type II model in an autoregressive framework**.

**S16 Table. Mean parameter estimates and credible intervals resulting from fitting the MFP, Type II model to five stochastic time series generated by the stable limit cycle parametrisation MFP, Type II model in an autoregressive framework**.

**S17 Table. Mean parameter estimates and credible intervals resulting from fitting the MFP, Type II model to five stochastic time series generated by the divergent oscillations parametrisation MFP, Type II model in an autoregressive framework**.

**S18 Table. Mean parameter estimates and credible intervals resulting from fitting the MFP, Type III model to five stochastic time series generated by the MFP, Type III model in an autoregressive framework**.

**S5 Fig. Stochastic time series and estimated population abundances over time**. After fitting generating models to “matching” stochastic time series in an autoregressive framework, we compared the stochastic time series to estimated population abundances. Each plot shows stochastic time series in solid lines, and estimated population abundances in dashed lines. Parasite population abundances are shown in green, while immune cell abundances are shown in blue. Labels on the left denote the generating model and time series. Cases in which the behaviour of the stochastic time series appears to be inconsistent with the estimated population abundances are highlighted by red boxes.

**S6 Fig. Overlaps in posterior distributions of consequential parameters**. Per the continuous-time implementation of the MFP, Type II model, differences in four parameters (*β, h, e*, and *δ*) produce damping oscillations, a stable limit cycle, or divergent oscillations. Each plot shows the posterior predictions that result from fitting the MFP, Type II model to different time series. In each plot, we show the posterior predictions on the vertical axis, and the specific, simulated time series on the horizontal axis. Upper and lower bounds denote the 95% credible interval. Black circles denote the true value. Grey shading indicates non-convergence.

**S7 Fig. Overlaps in posterior distributions of consequential parameters**. Per the discrete-time implementation of the MFP, Type II model, differences in two parameters (*r* and *δ*) produce damping oscillations, a stable limit cycle, or divergent oscillations. Each plot shows the posterior predictions that result from fitting the MFP, Type II model to different time series. In each plot, we show the posterior predictions on the vertical axis, and the specific, simulated time series on the horizontal axis. Upper and lower bounds denote the 95% credible interval. Black circles denote the true value.

